# Potentiation of EGFR mutant lung cancer treatment targeting replication stress

**DOI:** 10.1101/2025.04.18.649510

**Authors:** Reshma Bhowmick, John J. Turchi, Shadia I. Jalal

## Abstract

**Background/Objectives:** Oncogene-targeted therapies against growth factor receptors are effective treatment options for driver mutation-positive non-small cell lung cancer (NSCLC). The third-generation EGFR-Tyrosine Kinase inhibitor (TKI) osimertinib, is a standard first-line therapy for patients with EGFR-mutated NSCLC cancer. Acquired resistance to osimertinib is a significant problem that limits survival. In addition to that clinical data show approximately 15% of EGFR mutant non-small cell lung cancer patients have innate resistance to Osimertinib. Tumor heterogeneity and multiple resistance mechanisms add to the complexity of EGFR-mutated NSCLC. Tumor cells that acquire resistance, independent of the mechanism, experience replication stress (RS) as proliferation resumes.

**Methods:** We have employed a series of non-small cell lung cancer models to assess the efficacy of combining osimertinib with NERx-329, targeting RS via inhibiting RPA, and the impact of these treatments on growth and damage signaling pathways.

**Results:** We demonstrate that targeting RS with RPAi treatment induces cell death in multiple EGFR-mutant cell lines alone and when combined with osimertinib. Dissection of signaling pathways revealed that RPAi treatment does not block osimertinib inhibition of EGFR signaling. Analysis of DNA damage response (DDR) signaling reveals that NERx329 potentiates the osimertinib-dependent downregulation of Chk1 activity and expression. The Chk1 loss was shown to be dependent on proteasome degradation as proteasome inhibitor MG-132 restores Chk1 activity and expression.

**Conclusions:** From these data, we infer that targeting RS via RPAi NERx-329 in combination with osimertinib is an effective strategy and represents a promising drug combination targeted therapy to enhance efficacy and limit the development of resistance.

**Simple Summary:** This research aims to assess a new mechanism to potentiate osimertinib treatment of EGFR-mutated non-small cell lung cancer. Targeting replication stress with the small molecule Replication proteins A inhibitor (RPAi), NERx-329, induces cancer cell death and enhance the effectiveness of osimertinib. The findings suggest that combining NERx-329 with osimertinib could be a promising strategy to improve treatment outcomes and reduce resistance. Importantly, we demonstrate that neither RPAi or osimertinib alter the anticancer mechanism of each single agent, providing a new approach to increase therapeutic efficacy and delay or eliminate the development of drug resistance.

## 1. Introduction

The treatment of EGFR mutant non-small-cell lung cancer (NSCLC) harboring epidermal growth factor receptor (EGFR) activating mutations in exon 19 or 21 has significantly evolved in the last few years [1, 2]. EGFR tyrosine kinase inhibitors (TKIs) approved as single agents in the metastatic first line setting include osimertinib (the preferred option) in addition to Dacomitinib or Afatinib. Combination therapies have also been recently approved with EGFR TKIs forming the backbone of these regimens. EGFR-TKI’s are cytostatic and thus require continual treatment to maintain tumor control. However, prolonged use of these inhibitors results in resistance in most patients. While initial mechanisms of resistance were defined and effectively targeted with second and third generation TKI’s, many are unknown and therefore not effectively targeted [3, 4]. Moreover, multiple resistance mechanisms can arise in a single patient due to tumor heterogeneity which presents a significant clinical hurdle [5]. Effective mechanisms to combat TKI resistance in EGFR-driven NSCLC remain to be discovered.

Third-generation EGFR-TKI, osimertinib is a mutation specific EGFR inhibitor active against various EGFR mutations including T790M mutants but not wild-type EGFR. Moreover, osimertinib displays high selectivity with limited activity against other kinases including AXL, AKT1 or HER3 [6, 7]. Unfortunately, continuous osimertinib treatment necessitated by the mechanism of action results in the development of resistance by various mechanisms [8]. These mechanisms include the acquisition of the EGFR-C797S mutation, loss of the T790M mutation, activation of growth signaling bypass pathways, and histological transformation including transformation to small-cell lung cancer [9, 10]. More recently combination regimens have been approved for first line therapy of EGFR mutant NSCLC including osimertinib in combination with carboplatin and pemetrexed and the combination of Amivantamab with Lazertinib. It is unclear if these regimens are more effective than single agent osimertinib and are associated with increased toxicity [11, 12]. While the development of 4^th^ generation EGFR inhibitors is underway, alternatives to prolonged cytostatic activity could be beneficial [8].

Recent studies have identified alterations in the DNA damage response in osimertinib resistant cells and tumors suggestive that targeting DNA replication stress could be an effective strategy to limit the development of resistance and potentially overcome already established resistance [13-16]. Targeting resistant cells as they emerge would increase progression free survival and prolong the effectiveness of EGFR targeted TKIs. As resistant cells are defined by their re-entry into and progression through the cell cycle, these cells would be uniquely sensitive to agents targeting the replication stress and DNA damage response pathways. The DNA replication stress response is initiated by replication protein A (RPA) binding to the single strand DNA (ssDNA) intermediates emanating from replication stress. RPA binding initiates the ATR signaling pathway to limit replication, the generation of additional ssDNA and cellular exhaustion of RPA [17, 18]. Exhaustion of RPA can be accomplished genetically, chemically via small molecule inhibitors of the RPA-DNA interaction (RPAi) or directly via ATR inhibition [17, 19, 20]. In this study, we focus on combining RPAi treatment with osimertinib as a mechanism to induce cell death in EGFR-mutant NSCLC.

## 2. Materials and Methods

### Materials

osimertinib and Alectinib were obtained from Selleckchem (Houston, TX). Replication protein A inhibitor (NERx-329) was synthesized in our lab as previously described [19].

### Cell culture

Four Human NSCLC cell lines were employed in these studies (Table 1). The exon 19 deleted cell lines HCC4006 (L747-E749) and PC9 (E746-A750), the exon 21 double mutated H1975 cells (L858R/T790M), and the H2228 (EML4-ALK fusion gene) were obtained from commercial vendors. All cells were cultured in in RPMI-1640 media supplemented with 10% Fetal Bovine Serum (FBS) and 1xpennicilin/streptomycin under standard cell culture conditions.

**Table 1.**
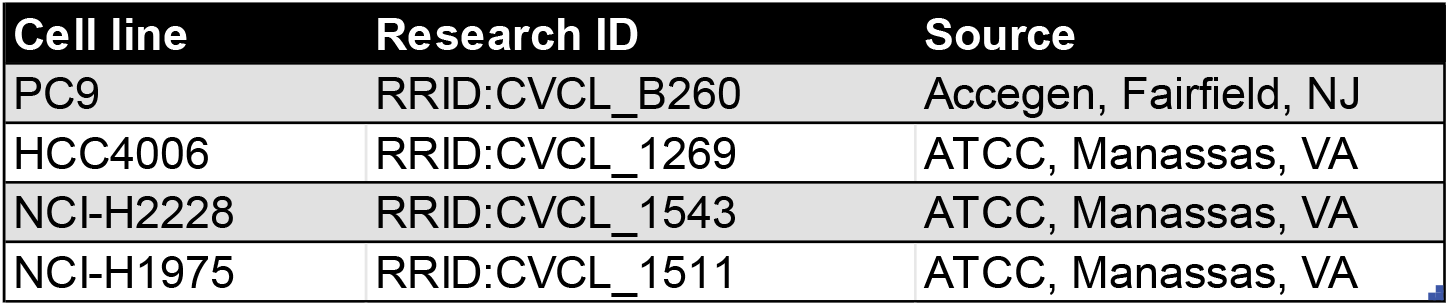
Cell lines used in the study.

| Cell line | Research ID | Source |
| --- | --- | --- |
| PC9 | RRID:CVCL_B260 | Accegen, Fairfield, NJ |
| HCC4006 | RRID:CVCL_1269 | ATCC, Manassas, VA |
| NCI-H2228 | RRID:CVCL_1543 | ATCC, Manassas, VA |
| NCI-H1975 | RRID:CVCL_1511 | ATCC, Manassas, VA |

### Metabolic cell viability assay

Cell viability is measured by metabolic CCK-8 assay (Dojindo laboratories) as we have described previously [20]. Briefly, tumor cells (2.5-5×10^3^ cells/well) were plated in 96-well plates and grown for 24h before adding osimertinib and NERx-329. Following this period of growth, cells were incubated with the indicated concentration of DMSO soluble compounds osimertinib and NERx-329 for 48h. Appropriate DMSO controls were used to compare with experimental samples. After 48h of incubation, CCK-8 reagent was added, and cells were incubated for 1-2 hours in a tissue culture CO_2_ incubator. Finally, the absorbance of formed formazan was measured at 450 nm using BioTek Synergy H1 Plate Reader (Agilent Tech, Santa Clara, CA). The percentage viability was determined relative to vehicle (DMSO) controls. The mean and SEM are plotted from triplicate determinations.

### Clonogenic survival assay

PC9 cells were plated in 6-well plates and following 24 h of growth, cells were treated with compounds. The final concentration of DMSO used for all treatments was 0.5%. The concentration of the compounds is indicated in **Fig. 2**. After 48h of treatment cells were trypsinized and equal numbers of cells were plated in 10cm dishes. Plated cells were incubated at 37°C in 5% CO2 for 11 days in a tissue culture incubator. The colony formation was detected by staining with 0.5% crystal violet supplemented with 6% glutaraldehyde in PBS. Colonies were manually counted, and the surviving fraction was normalized to vehicle controls.

### Incucyte Live Cell proliferation assay

The NSCLC cell lines were seeded in 96 well plates at 5×10^3^ cells/well. After 24h, the vehicle or indicated compounds were added and cell proliferation was monitored by Incucyte S3 Automated Live-Cell Analysis (Sartorius) for 7 days. Phase contrast images were collected every 4 hours and percentage confluence was determined. Data was presented as the mean and SEM of triplicate data points as we have described [20, 21].

### Antibodies and Western Blotting

The human NSCLC cells were plated in 6-well plates and grown for 18-24h. Cells were then treated with indicated concentrations of compounds and time points. The involvement of the proteasome in the degradation of chk1 was tested by adding proteasome inhibitor MG-132 (Cat: M7449, Sigma-Aldrich Co, St Louis, MO) to the cells along with other compounds or 12h later adding other compounds. Cell lysates were prepared in RIPA buffer (50 mM Tris, pH 8.0, 150 mM sodium chloride, 1% NP-40, 0.5% sodium deoxycholate, 0.1% SDS, 1mM EDTA) with protease and phosphatase inhibitors (Halt™ Protease and phosphatase inhibitor, single-use cocktail (100×). Cat: 78443, Thermo Scientific) by sonication.

The protein concentrations were determined by BCA assay (Pierce BCA protein assay kit, Thermo-scientific, Rockford, IL). Cell lysates of equal amounts (typically 50 mg) were resolved by SDS polyacrylamide gel electrophoresis (Bio-Rad, Hercules, CA) and transferred to polyvinylidene difluoride membranes (Cytiva, Marlborough, MA). The membranes were incubated with 5% w/v non-fat dry milk for 1h at room temperature then incubated with primary antibodies to p-Chk1, Chk1, p-EGFR, EGFR, p-ATR, ATR, p-Chk2, Chk2, and GAPDH following manufacturers (Cell Signaling Technology, Danvers, MA, USA,) at the manufacturer suggested dilution. Membranes were washed three times with and incubated with HRP-conjugated species-specific secondary antibody for 1h at room temperature. Finally, Clarity Western ECL Substrate (Biorad, Hercules, CA) was used to visualize the immunoreactive bands.

## 3. Results

### 3.1. NERx-329 – osimertinib combination in EGFR mutant NSCLC

#### 3.1.1. Mitochondrial assessment of viability

To determine the impact of combination RPAi -TKI therapy, we tested the effect of the RPAi NERx-329 on sensitization of three different EGFR-mutant NSCLC cell lines to osimertinib using CCK-8 metabolic assay after 48 hours of treatment (**Fig. 1)**. NERx-Dosedependent changes in cellular viability of HCC4006 cells in response to increasing concentrations of NERx-329 are shown in **Fig. 1A**. The IC_50_ of NERx-329 were determined to be 4.8 µM, which was reduced to 3.4 µM and 2.3 µM by combining with 2.5 nM and 5 nM osimertinib, respectively. To confirm this effect, we investigated osimertinib dosedependent changes on the viability of these cells in the presence of two different concentrations of NERx-329, the approximate IC_50_ and half the approximate IC_50_ **(Figure 1B)**. A similar reduction was observed in the osimertinib IC_50_ comparatively as the IC_50_ of 7.5 nM was reduced to 5.6 nM and 2.4 nM by adding 2.5 µM and 5 µM of NERx-329, respectively. To check the generality of this cytotoxic effect, we tested the PC-9 cell line another exon-19 deleted cell line. A similar potentiation of cytotoxicity was observed in these cells via compound combination as shown in the NERx-329 dose-response curve (**Fig. 1C**) and osimertinib dose-response curve (**Fig. 1D**). Although PC-9 cells showed a higher osimertinib IC_50_ of 46.8 nM compared to the HCC4006 cells, the addition of NERx-329 again revealed an increase in cytotoxicity. The H1975 NSCLC cell line revealed a similar sensitivity to RPAi treatment with an IC_50_ if ∼5 μM (**Fig. 1E**). The addition of osimertinib impacted cell viability and dramatically altered the dose response cure for RPAi treatment though accurate determination of IC_50_ values was hampered by the non-sigmoidal response to the combination treatment. The H1975 cells displayed considerably reduced sensitivity to osimertinib compared to the H4006 and PC9 cells, and a non-sigmoidal dose response (**Fig. 1F**) as has been previously reported [22]. The impact of subtoxic doses of RPAi was minimal in the H1975 cells. While interpretation of these shortterm viability experiments are limited in the H1975 cells, collectively these data demonstrate that NERx-329 can potentiate the activity of osimertinib in TKI-sensitive EGFR mutant cancer cell lines. This effect of NERx-329 appears to be specific to EGFR mutant cancers as the response of EML4-ALK driven H2228 NSCLC to Alectinib was not impacted by NERx-329 treatment. (**Suppl Fig. S1**).

**Figure 1.**
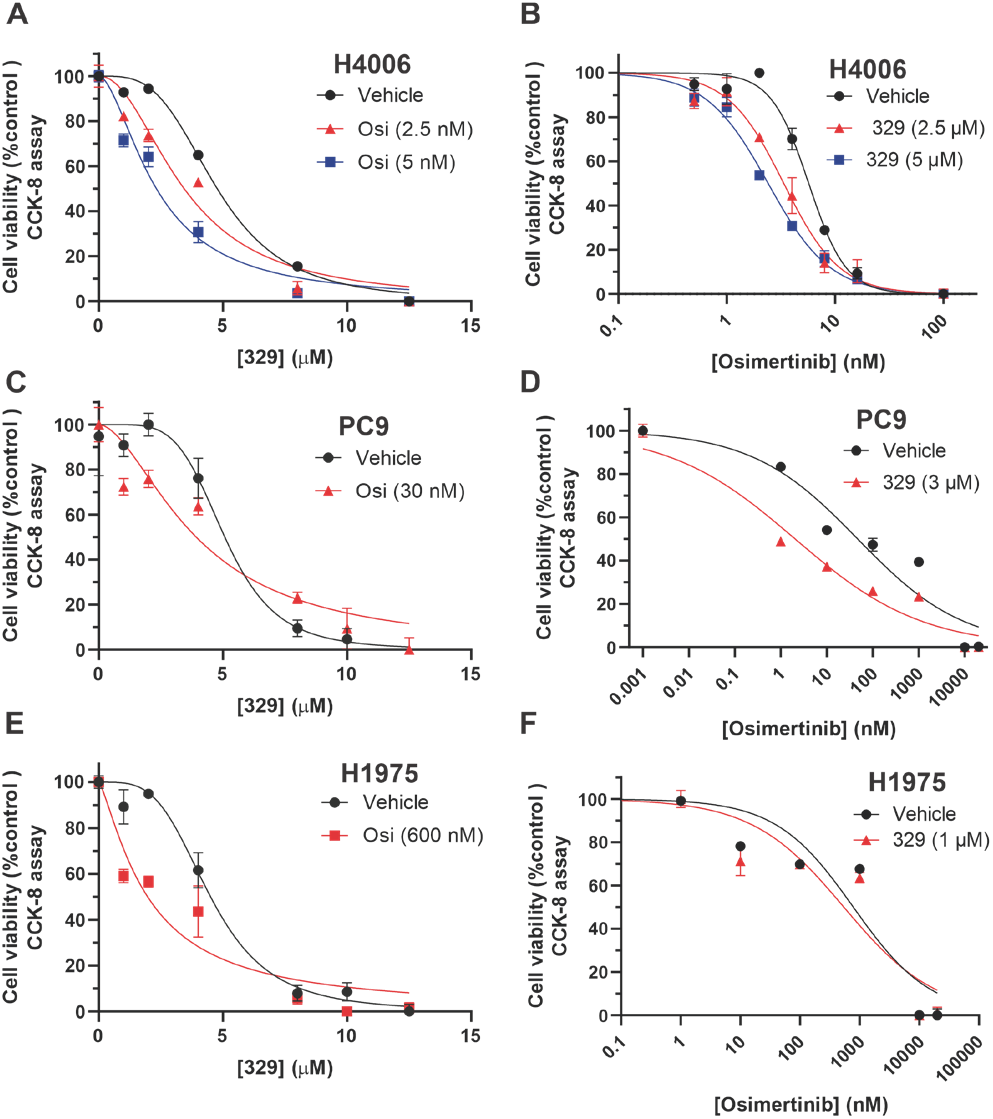
NERx329 potentiates the cytotoxicity of Osimertinib to NSCLC cell lines. The indicated EGFR-mutated NSCLCs were plated for 24h and then exposed to NERx-329 and Osimertinib at the indicated concentration for 48h. Cell viability was measured by metabolic CCK-8 assay. Data are presented as the mean and SEM of triplicate determinations.

#### 3.1.2. Clonogenic survival assessment of viability

The robust decrease in cell viability by the addition of NERx-329 with osimertinib prompted us to compare osimertinib alone clonogenic survival vs. combination treatment in the PC9 cell line cells (**Fig. 2**). The cells showed an osimertinib dose-dependent reduction in colony formation capacity which reduced to osimertinib (**Fig. 2A)**. Interestingly, adding the same amount of 329 reduces the cell survival of 5 nM osimertinib-treated cells from 33% to 9% (**Fig. 2A,B**). Similarly, adding NERx-329 reduces Osimertinib-induced cell survival at higher concentrations of Osimertinib (**Fig. 2B**).

**Figure 2.**
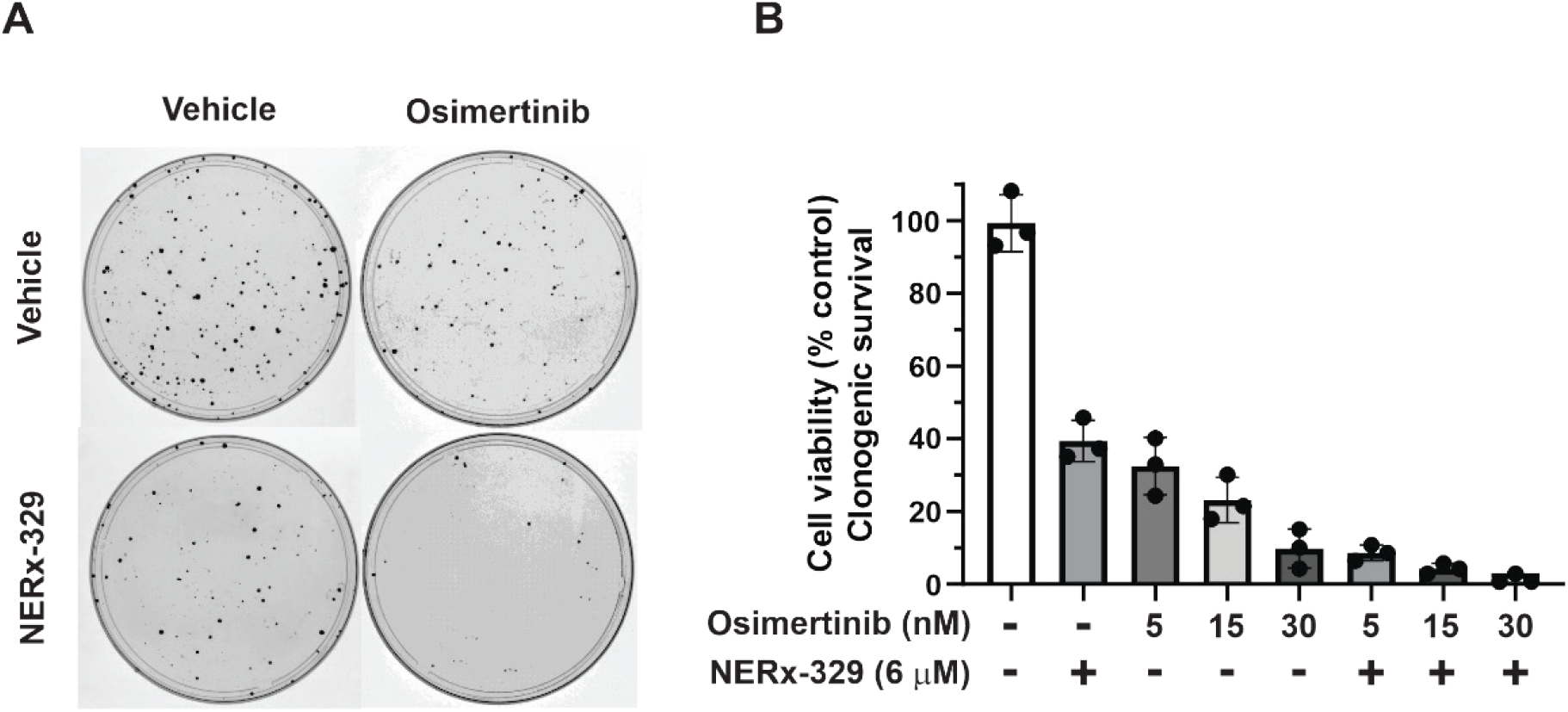
Clonogenic survival of PC9 NSCLC cells. (A) PC9 cells were treated with 5 nM Osimertinib and 6 μM NERx-329 as indicated, colonies were allowd to form over 11 days and were stained with crystal violet. (B) Quantification of the data presented as the mean, SD and individual values from triplicatae determiantions.

#### 3.1.3 Live cell imaging of NSCLC sensitivity to RPAi-TKI combination treatment

The reduction in cell viability as assessed by metabolic assays or clonogenic survival (**Figs. 1 and 2)** could result from the cytostatic activity of osimertinib or the cytotoxic effect of NERx-329. To gain more insight into the interactions and combined activity of osimertinib and NERx-329 we employed live cell imaging and extended treatment times of 7 to 8 days (**Fig. 3**). The extended time frame allowed identical concentrations to be used in the combinations. RPAi was employed at 2.5 μM and reduced proliferation by approximately 50% in each cell line, consistent with the tight range in which RPAi have been shown to be active [20]. Interestingly, 4 nM osimertinib was shown to be effective in reducing proliferation in each cell line despite the differences in sensitivity observed in the endpoint mitochondrial metabolic assay. The most significant impact on cell growth was observed after at least 48 hours of incubation with single-agent osimertinib, likely a result of the lag-time observed in achieving logarithmic growth. The RPAi-osimertinib combination resulted in a further significant reduction in proliferation compared to the single-agent treatments, in all cell lines tested. Interestingly, H1975 cells, which displayed resistance to osimertinib as assessed by the CCK-8 metabolic assay, exhibited markedly inhibited cell proliferation, via both osimertinib and NERx-329, over the 7-day experiment (**Fig. 3C**) and the combination treatment completely abrogated cell growth and resulted in a decrease in the normalized confluence below the initial normalized value.

**Figure 3.**
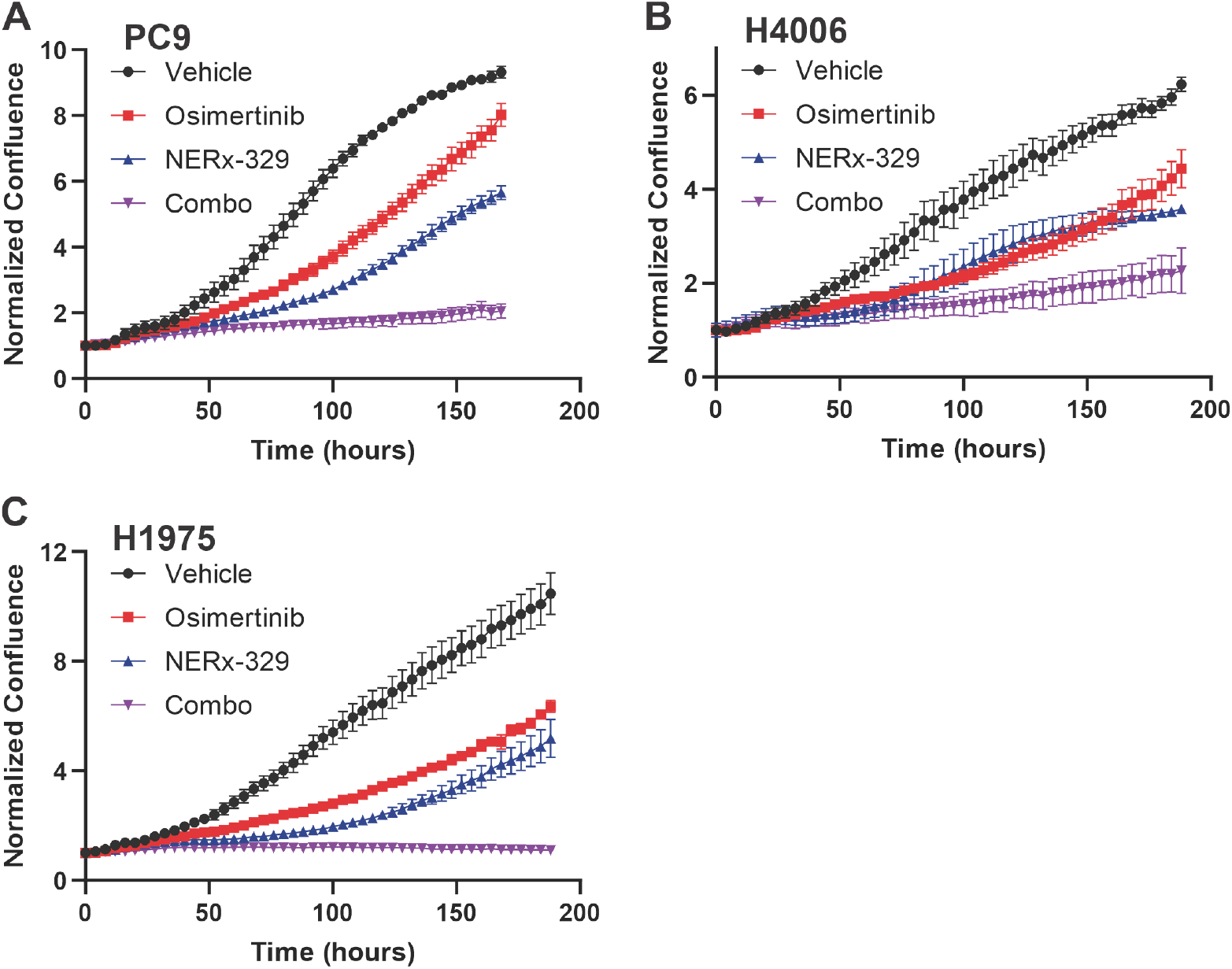
Live cell imagin of NSCLC cells treated with NERx-329 in combination with osimertinib. (A) PC-9, (B) HCC4006, and (C) H1975 cell lines were treated with 4 nM osimertinib and 2.5 µM NERx-329 as indicated. Images wer collected every 6 hours and normalized confluence determined. Data are plotted as the mean ± SEM of triplicate determinations.

### 3.2. osimertinib *inhibition of EGFR signaling is unaffected by RPAi treatment*

To elucidate the mechanism of combination activity we assessed the impact of RPAi treatment on osimertinib-dependent blockade of EGFR signaling. EGFR activation was assessed by the detection of phosphorylated EGFR at Tyrosine 1068 (Y1068) [7, 9]. PC-9 cells displayed a modest increase in total EGFR induced by NERx-329 at 6 hours. Osimertinib and combination treatment resulted in a dramatic increase in total EGFR expression 6 hours post treatment (**Fig. 4A**). These increases were still present but reduced after 24 hours of treatment. Activated EGFR as assessed by Y1068 phosphorylation is evident in untreated cells consistent with the driver mutation resulting in constitutive activation of the growth signaling pathway. Treatment with NERx-329 had a minimal effect on pEGFR levels while osimertinib resulted in a near complete reduction in p-EGFR level. The addition of NERx-329 did not negatively impact osimertinib-dependent blockade of EGFR phosphorylation observed at both the 6 and 24 h time points. Similar results were observed in the HCC4006 EGFR mutant NSCLC cell line (**Fig. 4B**). The NERx-329-dependent increase in total EGFR was less evident in these cells though the decrease in p-EGFR was stark in osimertinib treated cells independent of NERx-329 treatment.

**Figure 4.**
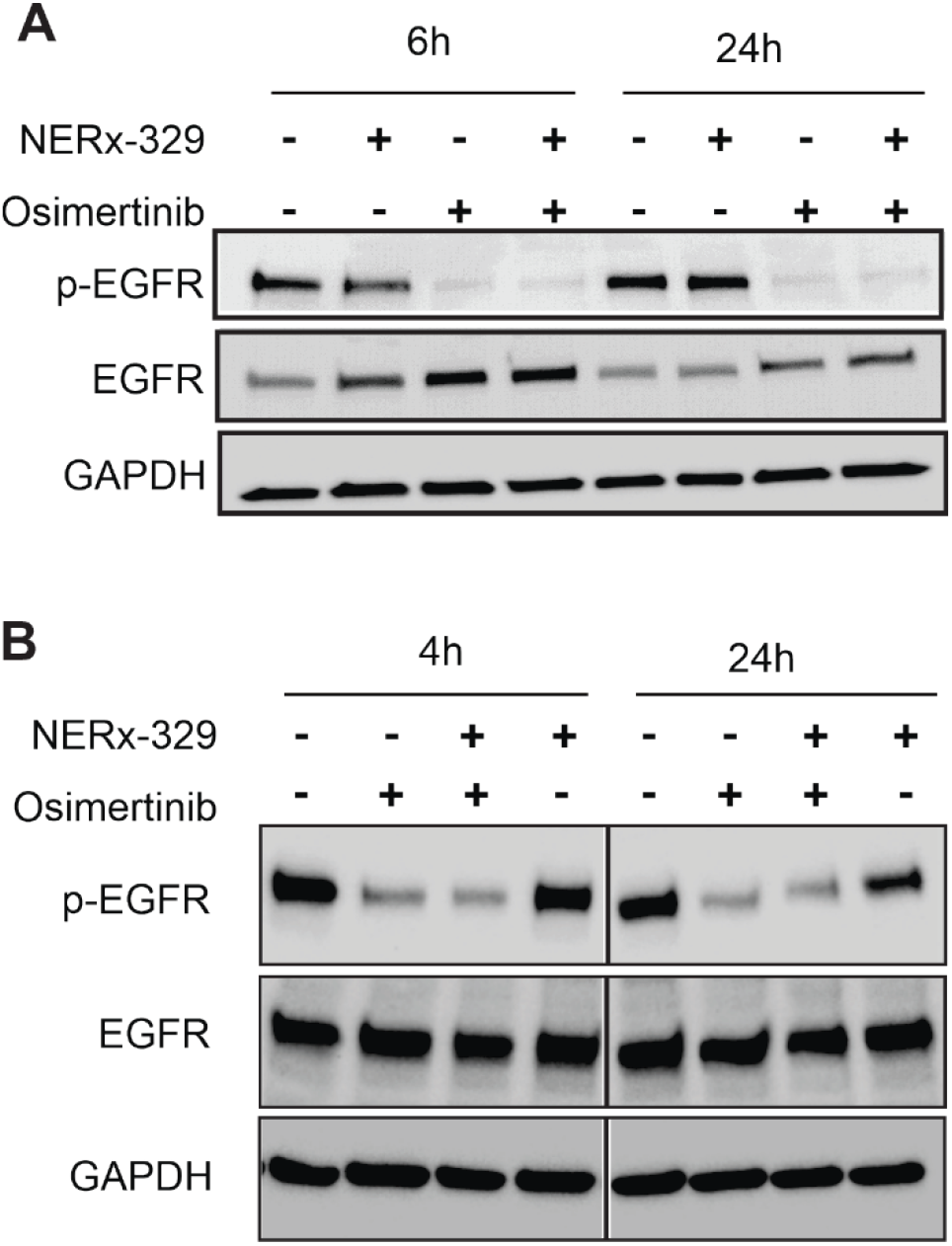
Impact of RPAi treatment on osimertinib-dependent EGFR blockade. (A) PC9 cells were treated with 30nM osimertinib and 1.5mM NERx-329 for the indicated times, cells were collected, lysed and loaded on SDS-PAGE and subjected to western blot analysis for p-EGFR, EGFR and GAPDH as described in “methods and Materials”. (B) HCC4006 cells were treated with 8 nM osimertinib and 5 mM NERx-329 as indicated and processed as described.

### 3.3 Impact of osimertinib on DDR signaling pathways

DDR signaling is impinged on by a variety of cellular events including DNA damage and replication stress. Dysregulated cell growth signaling pathways can induce replication stress by prematurely pushing cells into S-phase. Considering the effective combination treatment regimens (**Fig. 3**) we sought to determine how TKI treatment in EGFR mutant NSCLC cell lines impacted DDR signaling. A critical component of the DDR is the serine/threonine kinase Chk1. Chk1 is phosphorylated and activated by ATR in response to replication stress via excess single-stranded DNA initially sensed by RPA [23]. Interestingly, a low level of endogenous Ser345 phosphorylation was observed in PC9 and HCC4006 cells (**Fig. 5A, B**), consistent with endogenous replication stress. Analysis of PC9 cells treated with osimertinib and NERx-329 alone or in combination, revealed minimal changes in the level of Chk1 expression and activation as determined by detection of phosphorylation at Ser345 after 6h post-of treatment (**Fig. 5A**). Interestingly however, 24h following treatment we observed a significant reduction in the level of total Chk1 and a corresponding decrease of p-Chk1 in cells treated with osimertinib alone or in combination with NERx-329. Slightly different results were obtained in HCC4006 cells, with the magnitude of the reduction in Chk1 expression being less than that observed in PC-9 cells. Previous studies have demonstrated that in response to DNA damage, ATR-dependent Chk1 phosphorylation leads to Chk1 degradation [24].Consistent with this, is the low level of endogenous p-ATR observed in untreated PC-9 cells. Treatment with NERx-329, osimertinib, or the combination had no significant effect on the level of total ATR or its phosphorylation (**Fig. 5A**).

**Figure 5.**
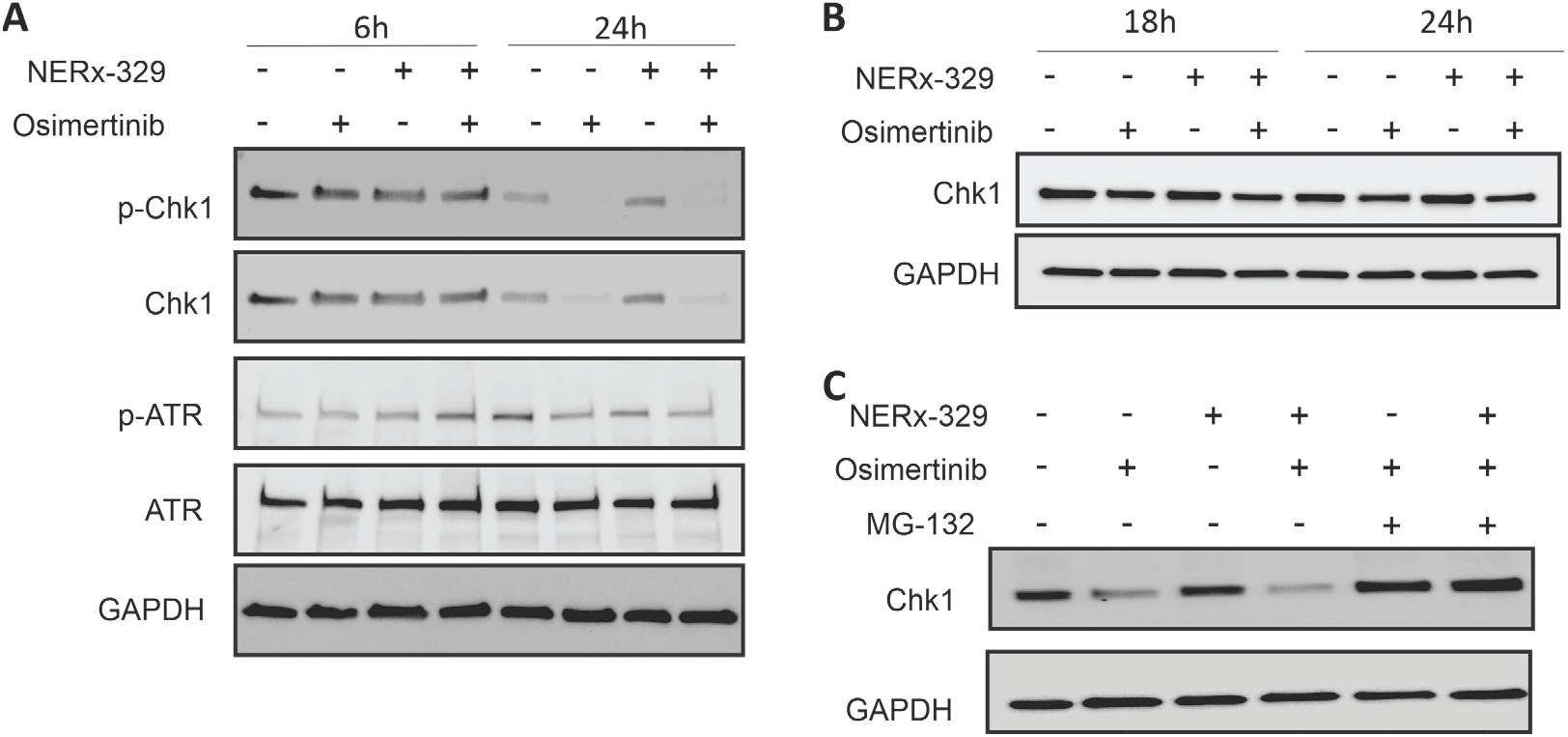
Impact of osimertinib on DDR signaling in unperturbed cells. (A) PC9 cells were treated with 30nM Osimertinib and 1.5 μM 329 as indicated and processed as described in Figure 4. (B) HCC4006 cells were treated with 4nM osimertinib and 2.5 μM NERx-329 as indicated. (C) PC9 cells were treated with 30 nM osimertinib, 2.5 μM NERx-329 and 2 μM MG-132 for 24 hours, as indicated. Cells were processed for western blot analysis of the indicated proteins as described above.

To determine the cause of osimertinib-dependent Chk1 degradation, we treated PC9 cells with osimertinib, NERx-329, and the combination in the presence of proteasomal inhibitor MG-132 (**Fig. 5C**). Treatment with osimertinib alone and in combination with NERx-329 showed decreased levels of Chk1 expression, as previously shown, and the addition of MG-132 abrogated the decrease in Chk1. These data demonstrate that TKI-dependent blockage of EGFR signaling results in proteosome-dependent Chk1 degradation. The data also demonstrates low-level, endogenous ATR activation and Chk-1 phosphorylation in unperturbed cells is insufficient to drive degradation of Chk1. Importantly, osimertinib-dependent proteosome-mediated Chk1 degradation is independent of RPA inhibition (**Fig. 5C**).

## 4. Discussion

In this study we have determined the impact of combining NERx-329, an RPA inhibitor, with osimertinib, a third-generation TKI, on EGFR-mutant NSCLC cell lines. The results demonstrate that NERx-329 enhances the cytotoxicity of osimertinib in EGFR-mutant cell lines, such as HCC4006 and PC-9. This potentiation effect is specific to EGFR-mutant cancers, as the combination did not affect EML4-ALK driven H2228 NSCLC cells treated with Alectinib. The study highlights the potential of NERx-329 to improve the efficacy of TKI-targeted therapy in EGFR-mutant lung cancer by increasing the sensitivity of cancer cells to osimertinib. The study also explored the mechanism behind enhanced cytotoxicity, showing that the combination treatment did not negatively impact the osimertinib-dependent blockade of EGFR phosphorylation. This suggests that NERx-329 enhances the therapeutic effect of osimertinib without interfering with its primary mechanism of action. We speculate that cell populations less sensitive to osimertinib that continue through the cell cycle and experience replication stress are sensitive to RPA inhibition. Additionally, the study examined the impact of the combination treatment on DDR signaling pathways. The results showed that osimertinib, alone or in combination with NERx-329, led to a significant reduction in Chk1 expression and activation in EGFR-mutant cell lines. This reduction was proteasome-dependent, as the addition of a proteasomal inhibitor abrogated the decrease in Chk1 levels. These findings suggest that the combination of NERx-329 and osimertinib not only enhances the cytostatic activity of TKI-targeted therapy but also impacts DDR signaling that can be exploited by DDR targeted agents. There is potential that the efficacy of the combination will reduce the development of resistance often observed in TKI treated EGFR-mutant NSCLC. We posit that cells which escape TKI-dependent inhibition of proliferation will be sensitive to RPAi inhibition and thus reduce the likelihood of resistance. EGFR mutant cells have been shown to be sensitive to other DNA damaging agents beyond platinum-based chemotherapy including PARP inhibitors [25]. EGFR mutant cells regardless of mechanism of acquired resistance to EGFR TKIs display increased sensitivity to PARP inhibitors when compared to their EGFR TKI sensitive counterparts [26]. The exact mechanisms through which EGFR TKI resistance induces PARP inhibitor sensitivity have not been fully defined but current data suggests this is not dependent on PARP trapping or its role in DNA repair. EGFR TKI resistant models all demonstrated a slight increase in reactive oxygen species at baseline which was further accentuated in response to PARP inhibitors. This was not observed in EGFR TKI sensitive models. PARP inhibitor sensitivity was reversed by antioxidants known to act as ROS scavengers in these resistant models. PARP inhibitor sensitivity was noted in “drug tolerant persister cells” which initially represent a small subpopulation in most patients but eventually expand playing a role in disease relapse, progression and the development of resistance. This data along with our work suggests DNA repair inhibitors can be exploited in EGFR mutant NSCLC.

## 5. Conclusions

This study has demonstrated the utility of combining the novel DDR targeted agent NERx-329 with osimertinib to treat EGFR-mutant NSCLC. The ability to induce cell death with a DDR targeted agent in combination with the cytostatic activity of TKI treatment offers the potential to allow altered dosing and treatment regimens to maximize efficacy and reduce the development of TKI resistance. The ability to maintain the mechanistic blocks of each agent in the combination therapy also suggests the combination can be delivered safely. While TKIs has revolutionized the treatment of EGFR-mutant NSCLC, there are few if any cures and new approaches are needed to overcome this clinical barrier. Thus, extensive preclinical analyses are needed to pursue these studies clinically. The results presented here demonstrate that RPAi treatment does not negatively impact TKI-dependent EGFR blockade, and that combination treatment offers increased cancer control. This represents an important step forward to developing novel therapeutic treatment regimens for EGFR-mutant NSCLC.

## Supplementary Materials

The following supporting information can be downloaded at: www.mdpi.com/xxx/s1, Figure S1:

### Author Contributions

Conceptualization, SJ and JT; methodology, SJ and JT; formal analysis, RB, SJ and JT.; investigation, RB; resources, SJ and JT; writing—original draft preparation, SB; writing— review and editing, SJ and JT; supervision, SJ; project administration, SJ; funding acquisition, SJ and JT. All authors have read and agreed to the published version of the manuscript

### Funding

This research was funded by the National Institutes of Health, grant number R01CA257430 and the Tom and Julie Wood Family Foundation (JJT). The APC was funded with support from the Tom and Julie Wood Family Foundation.

### Data Availability Statement

The original contributions presented in this study are included in the article/supplementary material. Further inquiries can be directed to the corresponding author(s).

## Acknowledgments

We thank all member of the Turchi Lab and NERx Biosciences for their scientific input and critical reading of the manuscript.

## Abbreviations

The following abbreviations are used in this manuscript:

EGFR: Epidermal growth factor receptor
RPA: Replication protein A
TKI: Tyrosine kinase inhibitor
RS: Replication stress
DDR: DNA damage response
NSCLC: Non-small cell lung cancer
ssDNA: Single-strand DNA

## Disclaimer/Publisher’s Note

The statements, opinions and data contained in all publications are solely those of the individual author(s) and contributor(s) and not of MDPI and/or the editor(s). MDPI and/or the editor(s) disclaim responsibility for any injury to people or property resulting from any ideas, methods, instructions or products referred to in the content.

## Reference List

1. Sha C, Lee PC: EGFR-Targeted Therapies: A Literature Review. J Clin Med 2024, 13.

2. Janne PA, Yang JC, Kim DW, Planchard D, Ohe Y, Ramalingam SS, Ahn MJ, Kim SW, Su WC, Horn L, Haggstrom D, Felip E, Kim JH, Frewer P, Cantarini M, Brown KH, Dickinson PA, Ghiorghiu S, Ranson M: AZD9291 in EGFR inhibitor-resistant non-small-cell lung cancer. N Engl J Med 2015, 372:1689–1699.

3. Dickerson H, Diab A, Al MO: Epidermal Growth Factor Receptor Tyrosine Kinase Inhibitors in Cancer: Current Use and Future Prospects. Int J Mol Sci 2024, 25.

4. Sequist LV, Waltman BA, Dias-Santagata D, Digumarthy S, Turke AB, Fidias P, Bergethon K, Shaw AT, Gettinger S, Cosper AK, Akhavanfard S, Heist RS, Temel J, Christensen JG, Wain JC, Lynch TJ, Vernovsky K, Mark EJ, Lanuti M, Iafrate AJ, Mino-Kenudson M, Engelman JA: Genotypic and histological evolution of lung cancers acquiring resistance to EGFR inhibitors. Sci Transl Med 2011, 3:75ra26.

5. Niederst MJ, Sequist LV, Poirier JT, Mermel CH, Lockerman EL, Garcia AR, Katayama R, Costa C, Ross KN, Moran T, Howe E, Fulton LE, Mulvey HE, Bernardo LA, Mohamoud F, Miyoshi N, VanderLaan PA, Costa DB, Janne PA, Borger DR, Ramaswamy S, Shioda T, Iafrate AJ, Getz G, Rudin CM, Mino-Kenudson M, Engelman JA: RB loss in resistant EGFR mutant lung adenocarcinomas that transform to small-cell lung cancer. Nat Commun 2015, 6:6377.

6. Finlay MR, Anderton M, Ashton S, Ballard P, Bethel PA, Box MR, Bradbury RH, Brown SJ, Butterworth S, Campbell A, Chorley C, Colclough N, Cross DA, Currie GS, Grist M, Hassall L, Hill GB, James D, James M, Kemmitt P, Klinowska T, Lamont G, Lamont SG, Martin N, McFarland HL, Mellor MJ, Orme JP, Perkins D, Perkins P, Richmond G, Smith P, Ward RA, Waring MJ, Whittaker D, Wells S, Wrigley GL: Discovery of a potent and selective EGFR inhibitor (AZD9291) of both sensitizing and T790M resistance mutations that spares the wild type form of the receptor. J Med Chem 2014, 57:8249–8267.

7. Cross DA, Ashton SE, Ghiorghiu S, Eberlein C, Nebhan CA, Spitzler PJ, Orme JP, Finlay MR, Ward RA, Mellor MJ, Hughes G, Rahi A, Jacobs VN, Red BM, Ichihara E, Sun J, Jin H, Ballard P, Al-Kadhimi K, Rowlinson R, Klinowska T, Richmond GH, Cantarini M, Kim DW, Ranson MR, Pao W: AZD9291, an irreversible EGFR TKI, overcomes T790M-mediated resistance to EGFR inhibitors in lung cancer. Cancer Discov 2014, 4:1046–1061.

8. Corvaja C, Passaro A, Attili I, Aliaga PT, Spitaleri G, Signore ED, de MF: Advancements in fourth-generation EGFR TKIs in EGFR-mutant NSCLC: Bridging biological insights and therapeutic development. Cancer Treat Rev 2024, 130:102824.

9. Finlay MRV, Barton P, Bickerton S, Bista M, Colclough N, Cross DAE, Evans L, Floc’h N, Gregson C, Guerot CM, Hargreaves D, Kang X, Lenz EM, Li X, Liu Y, Lorthioir O, Martin MJ, McKerrecher D, McWhirter C, O’Neill D, Orme JP, Mosallanejad A, Rahi A, Smith PD, Talbot V, Ward RA, Wrigley G, Wylot M, Xue L, Yao T, Ye Y, Zhao X: Potent and Selective Inhibitors of the Epidermal Growth Factor Receptor to Overcome C797S-Mediated Resistance. J Med Chem 2021, 64:13704–13718.

10. Leonetti A, Minari R, Mazzaschi G, Gnetti L, La MS, Alfieri R, Campanini N, Verze M, Olivani A, Ventura L, Tiseo M: Small Cell Lung Cancer Transformation as a Resistance Mechanism to Osimertinib in Epidermal Growth Factor Receptor-Mutated Lung Adenocarcinoma: Case Report and Literature Review. Front Oncol 2021, 11:642190.

11. Planchard D, Janne PA, Cheng Y, Yang JC, Yanagitani N, Kim SW, Sugawara S, Yu Y, Fan Y, Geater SL, Laktionov K, Lee CK, Valdiviezo N, Ahmed S, Maurel JM, Andrasina I, Goldman J, Ghiorghiu D, Rukazenkov Y, Todd A, Kobayashi K: Osimertinib with or without Chemotherapy in EGFR-Mutated Advanced NSCLC. N Engl J Med 2023, 389:1935–1948.

12. Cho BC, Lu S, Felip E, Spira AI, Girard N, Lee JS, Lee SH, Ostapenko Y, Danchaivijitr P, Liu B, Alip A, Korbenfeld E, Mourao DJ, Besse B, Lee KH, Xiong H, How SH, Cheng Y, Chang GC, Yoshioka H, Yang JC, Thomas M, Nguyen D, Ou SI, Mukhedkar S, Prabhash K, D’Arcangelo M, Alatorre-Alexander J, Vazquez Limon JC, Alves S, Stroyakovskiy D, Peregudova M, Sendur MAN, Yazici O, Califano R, Gutierrez C V, de MF, Passaro A, Kim SW, Gadgeel SM, Xie J, Sun T, Martinez M, Ennis M, Fennema E, Daksh M, Millington D, Leconte I, Iwasawa R, Lorenzini P, Baig M, Shah S, Bauml JM, Shreeve SM, Sethi S, Knoblauch RE, Hayashi H: Amivantamab plus Lazertinib in Previously Untreated EGFR-Mutated Advanced NSCLC. N Engl J Med 2024, 391:1486–1498.

13. Amato L, Omodei D, De RC, Ariano A, Capaldo S, Tufano CC, Buono R, Terlizzi C, Nardelli A, Del V, V, Palumbo R, Tuccillo C, Morgillo F, Papaccio F, Tirino V, Iommelli F, Della Corte CM, De R, V: Combined Therapeutic Strategies Based on the Inhibition of Non-Oncogene Addiction to Improve Tumor Response in. Cancers (Basel) 2024, 16.

14. Ko JC, Chen JC, Huang CH, Chen PJ, Chang QZ, Mu BC, Chen JJ, Tai TY, Suzuki K, Wang YX, Lin YW: Downregulation of Rad51 Expression and Activity Potentiates the Cytotoxic Effect of Osimertinib in Human Non-Small Cell Lung Cancer Cells. Chemotherapy 2024:1–14.

15. De RC, De R V, Tuccillo C, Tirino V, Amato L, Papaccio F, Ciardiello D, Napolitano S, Martini G, Ciardiello F, Morgillo F, Iommelli F, Della Corte CM: ITGB1 and DDR activation as novel mediators in acquired resistance to osimertinib and MEK inhibitors in EGFR-mutant NSCLC. Sci Rep 2024, 14:500.

16. Liang XM, Qin Q, Liu BN, Li XQ, Zeng LL, Wang J, Kong LP, Zhong DS, Sun LL: Targeting DNA-PK overcomes acquired resistance to third-generation EGFR-TKI osimertinib in non-small-cell lung cancer. Acta Pharmacol Sin 2021, 42:648–654.

17. Toledo LI, Altmeyer M, Rask MB, Lukas C, Larsen DH, Povlsen LK, Bekker-Jensen S, Mailand N, Bartek J, Lukas J: ATR prohibits replication catastrophe by preventing global exhaustion of RPA. Cell 2013, 155:1088–1103.

18. Toledo L, Neelsen KJ, Lukas J: Replication Catastrophe: When a Checkpoint Fails because of Exhaustion. Mol Cell 2017, 66:735–749.

19. Gavande NS, VanderVere-Carozza PS, Pawelczak KS, Vernon TL, Jordan MR, Turchi JJ: Structure-Guided Optimization of Replication Protein A (RPA)-DNA Interaction Inhibitors. ACS Med Chem Lett 2020, 11:1118–1124.

20. VanderVere-Carozza PS, Gavande NS, Jalal SI, Pollok KE, Ekinci E, Heyza J, Patrick SM, Masters A, Turchi JJ, Pawelczak KS: In Vivo Targeting Replication Protein A for Cancer Therapy. Front Oncol 2022, 12:826655.

21. Mendoza-Munoz PL, Gavande NS, VanderVere-Carozza PS, Pawelczak KS, Dynlacht JR, Garrett JE, Turchi JJ: Ku-DNA binding inhibitors modulate the DNA damage response in response to DNA double-strand breaks. NAR Cancer 2023, 5:zcad003.

22. Tang ZH, Jiang XM, Guo X, Fong CM, Chen X, Lu JJ: Characterization of osimertinib (AZD9291)-resistant non-small cell lung cancer NCI-H1975/OSIR cell line. Oncotarget 2016, 7:81598–81610.

23. Zou L, Elledge SJ: Sensing DNA damage through ATRIP recognition of RPA-ssDNA complexes. Science 2003, 300:1542–1548.

24. Zhang YW, Otterness DM, Chiang GG, Xie W, Liu YC, Mercurio F, Abraham RT: Genotoxic stress targets human Chk1 for degradation by the ubiquitin-proteasome pathway. Mol Cell 2005, 19:607–618.

25. Pfaffle HN, Wang M, Gheorghiu L, Ferraiolo N, Greninger P, Borgmann K, Settleman J, Benes CH, Sequist LV, Zou L, Willers H: EGFR-activating mutations correlate with a Fanconi anemia-like cellular phenotype that includes PARP inhibitor sensitivity. Cancer Res 2013, 73:6254–6263.

26. Marcar L, Bardhan K, Gheorghiu L, Dinkelborg P, Pfaffle H, Liu Q, Wang M, Piotrowska Z, Sequist LV, Borgmann K, Settleman JE, Engelman JA, Hata AN, Willers H: Acquired Resistance of EGFR-Mutated Lung Cancer to Tyrosine Kinase Inhibitor Treatment Promotes PARP Inhibitor Sensitivity. Cell Rep 2019, 27:3422–3432.

