## Supplemental Figure 1 for "Potentiation of EGFR mutant lung cancer treatment targeting replication stress"

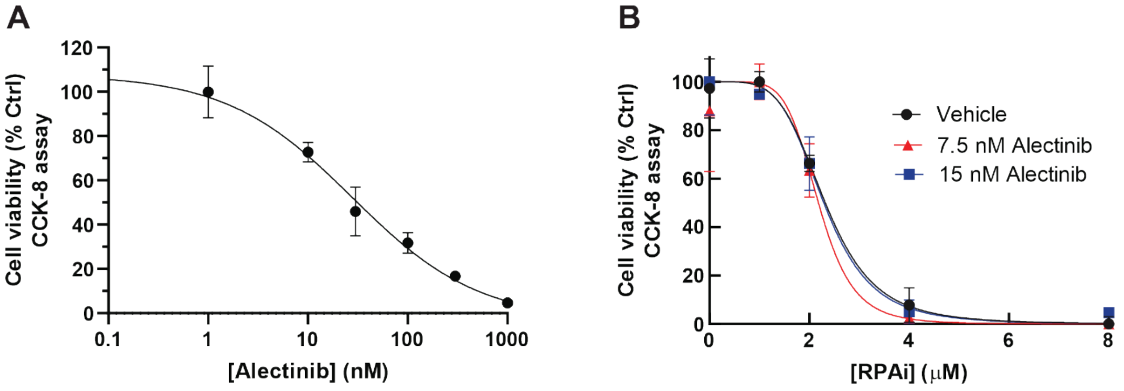


**Figure S1**. NERx-329 does not potentiate cytotoxicity of ALK TKI Alectinib in H2228-Alk driven NSCLC cells. (A) H2228 NSCLC cells were plated in a 96-well plate and grown for 24h. Cells were treated for 48 hours with the indicated concentration of Alectinib and cell viability assessed by CCK-8 assay (B) H2228 cells were plated as described above and Alectinib, 7.5 and 15 nM was added concurrent with the indicated concentrations of RPAi NERx-329. Cell viability was measured by metabolic CCK-8 assay 48-hours after treatment. Means and SEM are presented from triplicate determinations.
